# Molecular atlas of key food odorants reveals structured aroma organization and enables generative aroma design

**DOI:** 10.64898/2026.01.21.700072

**Authors:** Jingzhi Zhang, Huadong Xing, Antonella Di Pizio, Qinfei Ke, Xingran Kou, Dachuan Zhang

## Abstract

Food aromas arise from complex combinations of odorants, yet how these combinations are organized across foods to define aroma identity remains unclear. Decoding this “aroma code” could help bridge the sensory gap between traditional foods and sustainable alternatives, which often struggle with off-notes or to replicate consumer-expected flavor profiles. Here we present KFO-Atlas, a molecular atlas of 896 key food odorants curated from 2,282 food aroma formulations. Analysis shows that food aromas are built from sparse and structured sets of odorants. Plant-derived foods span a broad and diverse aroma space, whereas animal-derived foods tend to exhibit more similar odorant sets. In specific cases, distinct plant- and animal-based foods converge on similar odorant compositions through shared reaction pathways. Building on these insights, we develop a generative AI model that produces category-targeted aroma formulations and validate its outputs by blinded human sensory evaluation. As a proof of principle, the model reconstructs meat-like aromas using exclusively plant-derived odorants, demonstrating a data-driven route to address sensory bottlenecks in sustainable food products.

## Introduction

Aroma is a primary sensory signal of food, shaping first impressions, eating decisions, and consumer acceptance^1–3^. Food aroma arises from context-dependent combinations of odorants shaped by diverse metabolic origins and food processing. As a result, it remains difficult to systematically compare food aromas or to identify general principles that link odorant combinations to stable and recognizable aroma identity across foods^4,5^. This hinders both the mechanistic understanding and translation into controllable formulations.

Although over 10,000 volatile compounds have been identified in foods, key food odorants (KFOs), defined as volatile compounds with odor activity values greater than one, exert a disproportionate influence on aroma perception^5,6^. Early compilations estimated that only a small fraction of volatiles—approximately 230 (less than 3%)—qualify as KFOs^7^. This estimate likely underrepresents the true diversity of odor-active compounds, given the substantial advances in analytical sensitivity over the past decade. As the food aroma literature has expanded rapidly across diverse matrices and processing contexts, KFO data have accumulated in a fragmented manner across studies^5,8^. Consequently, the field lacks a comprehensive representation of the KFO landscape that would allow systematic comparison across foods and a mechanistic understanding of how food aroma is organized.

This gap also constrains aroma design. Despite the recognition of KFOs, food aroma formulation remains largely empirical because aroma perception arises from nonlinear interactions among multiple odorants^9^. The resulting combinatorial space is vast and cannot be exhaustively explored experimentally^10^. Existing computational approaches have largely focused on predicting sensory attributes of individual odorants^11–13^ or have been restricted to a few specific product categories, such as wine or whisky^14,15^, where sufficient data are available. As a result, the field lacks a general framework that links the KFOs to controllable aroma formulation.

Here, we introduce a conceptual and methodological framework to decode the molecular organization of food aroma. We construct KFO-Atlas, a comprehensive database of 896 KFOs, spanning diverse food categories. Using this resource, we systematically examine how KFO combinations are organized across foods, and whether common structural patterns underpin aroma identity and similarity. In particular, we ask whether the frequency with which individual KFOs appear across foods reflects their functional roles in shaping aroma identity. We further explore the extent to which distinct plant- and animal-based foods may converge on similar KFO composition under shared reaction pathways. Building on this organization, we develop a conditional variational autoencoder to test whether structured knowledge of KFO combinations can be translated into controllable aroma formulation. We validate the perceptual category fidelity of the generated formulations through a blinded sensory evaluation, in which experienced panelists assessed 100 AI-generated formulations for their perceived aroma category. As a proof of principle, we further apply the framework to generate meat-like aromas using exclusively plant-derived odorants.

## Results

### Overview of the key food odorants in KFO-Atlas

KFO-Atlas compiles 2,282 food aroma formulations curated from 1,667 publications published between 2015 and 2024, covering 20 food categories and 87 subcategories based on FoodEx2^16^ classification (**Supporting Data 1**). Across all formulations, we identified 896 unique KFOs, with the central 95% of odor activity values spanning 1.05 to 9,332.54. Despite this broad chemical repertoire, individual formulations relied on only a limited subset of KFOs: the central 95% contained between 3 and 33 odorants (**Figure 1a**). This narrow range indicates that food aroma identity is encoded by sparse combinations of KFOs.

**Figure 1.**
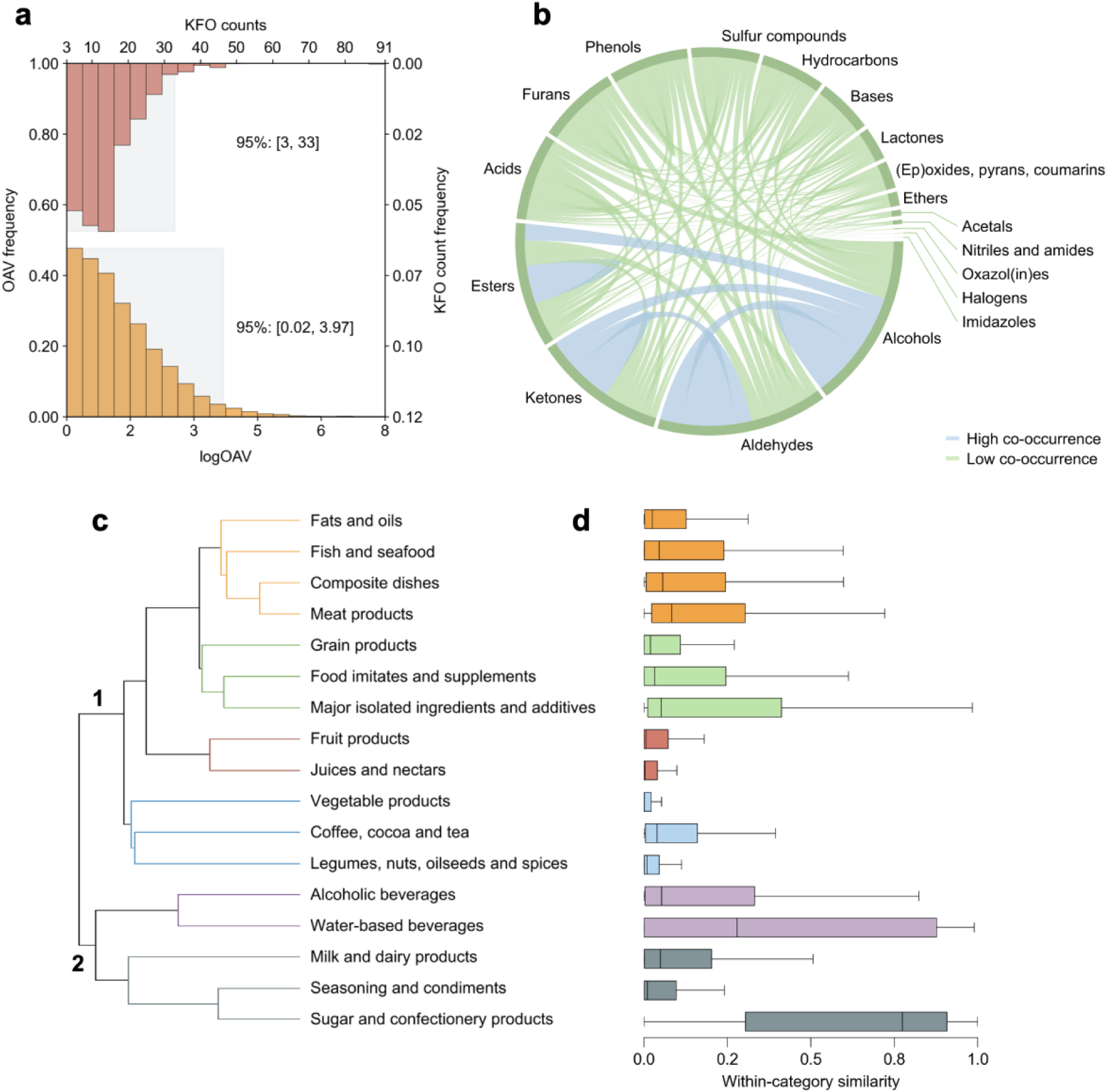
Global organization of key odorants across foods. Food aromas are built from sparse combinations of key odorants that give rise to structured organization across chemical classes and food categories. (a) Distribution of the number of KFOs per food aroma formulation and the corresponding odor activity values (OAVs) of individual KFOs. The blue contour encloses the central 95% of observations. (b) Network representation of chemical co-occurrence across food aroma formulations, in which nodes represent KFO chemical groups and edges denote co-occurrence frequency. The width of the outer arcs reflects the total co-occurrence of each chemical group across formulations. Edge colors indicate co-occurrence frequency, categorized using a threshold of 1,000 joint occurrences to distinguish high- and low-frequency co-occurrences. (c) Hierarchical clustering of 17 major food categories with at least ten formulations based on the cross-category similarity of their KFO compositions, showing that foods organize into distinct groups in KFO composition space. (d) Distributions of within-category pairwise similarity scores among KFO formulations for each food category. Lower similarity scores indicate greater KFO compositional heterogeneity, whereas higher scores reflect stronger convergence toward shared KFO compositions. Boxplots display the median and interquartile range, with whiskers extending to 1.5× interquartile range.

KFOs were classified into 18 chemical groups based on molecular structure (see *Methods*). Esters (20%), alcohols (13%), aldehydes (11%), ketones (11%), sulfur compounds (10%), and hydrocarbons (9%) accounted for most identified KFOs (**Table S1**). However, chemical abundance alone did not fully explain their roles in aroma organization. In the co-occurrence network, alcohols and aldehydes emerged as the most connected nodes, indicating they may act as connectors between diverse aroma compositions (**Figure 1b**). Consistent with this, most co-occurrence relationships (71%) occurred between different chemical classes rather than within the same class, suggesting that food aromas are built from mixtures of chemically distinct odorants rather than from single-class combinations.

The number of KFOs per formulation varied across food categories (**Table S2**). Categories such as coffee, cocoa, and tea (mean 16.49), seasonings and condiments (17.05), and legumes, nuts, oilseeds, and spices (17.02) showed the highest KFO counts, whereas water-based beverages (9.36), fish and seafood (10.63), and dairy products (11.29) exhibited much lower numbers of KFO counts. Collectively, these analyses establish a quantitative baseline for comparing aroma composition across foods and for linking sparse KFO sets to aroma identity.

### Structured aroma organization revealed by similarity and clustering analyses

To assess variation in aroma composition within food categories, we computed pairwise similarity scores among the KFO composition of food samples within each category (Figure 1d; see *Methods*). Lower within-category similarity scores indicate higher heterogeneity among KFO compositions, whereas higher scores reflect convergence toward shared compositions. Plant-derived categories, including vegetables, fruits, juices and nectars, and coffee, cocoa and tea, exhibited the lowest similarity. In contrast, animal-derived categories showed higher similarity, despite often containing larger numbers of KFOs (**Table S2**).

Hierarchical clustering based on cross-category KFO composition similarity further organized food categories into distinct groups (Figure 1c). Cluster 1 comprised categories associated with either diverse biosynthetic origins or thermally and lipid-driven chemistry. Within this cluster, plant-based categories occupied a broad region of the aroma space, accompanied by a broad distribution of alcohols, aldehydes, and other biosynthetically derived structural classes (**Figure S1**). By comparison, animal-derived foods formed tight subclusters characterized by highly similar KFO compositions.

Cluster 2 comprised food categories with relatively more similar KFO compositions. Water-based beverages contained the fewest KFOs and showed high within-category similarity, reflecting chemically simple matrices with limited fermentation or thermal transformation. Alcoholic beverages, although comprising many fermentation-derived KFOs, also exhibited strong convergence across samples. Milk and dairy products, sugar and confectionery products likewise showed high similarity, yielding stable chemical profiles despite differences in raw materials and processing (**Figure S1**). These clusters highlight marked differences in compositional heterogeneity across food categories, with some foods exhibiting highly diverse KFO profiles, whereas others show a relatively constrained and similar set of key odorants.

### Frequency-based hierarchy of key odorants shapes food aroma identity

To characterize how key odorants are distributed across foods, we classified all 896 KFOs based on their frequency across food aroma formulations into generalists (present in >25% of samples), intermediaries (5-25%), and individualists (<5%) following the previous approach^7^ (Figure 2a). Only six compounds—2-phenylethanol, linalool, 1-octen-3-ol, ethyl hexanoate, 1-hexanal, and nonanal—qualified as generalists and were detected across most food categories. Intermediaries (n=62) show moderate prevalence and typically arise from pathway activations tied to botanical origin, fermentation, or thermal processing. The vast majority (828; 92.4%) were individualists, including 164 odorants found only once.

**Figure 2.**
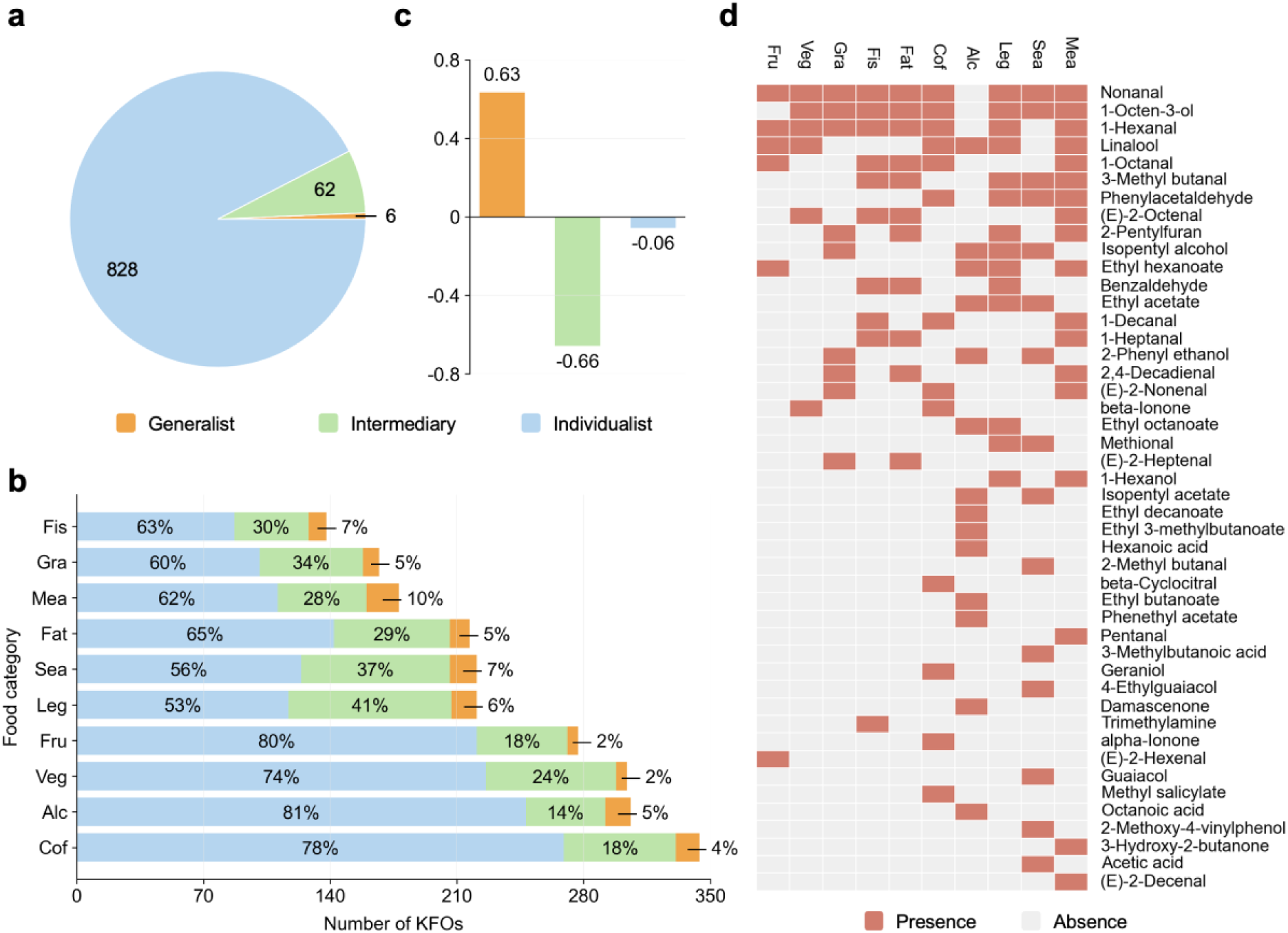
Frequency-based organization of key food odorants (KFOs) across food categories. (a) Distribution of 896 KFOs classified into three frequency-based groups across 2,282 food aroma formulations: generalists (>25%), intermediaries (5-25%), and individualists (<5%). (b) Counts of generalists, intermediaries, and individualists identified within each of the ten balanced food categories analyzed in this study. (c) Correlations between the abundance of each KFO frequency group within a food category and its within-category compositional similarity score. Spearman correlation coefficients (r) are shown for each group. (d) Presence-absence matrix of all generalist KFOs across food categories, illustrating their distribution across categories and the occurrence of category-specific generalists. Note: In panels (b) and (d), food category abbreviations are as follows: fis, fish and seafood; gra, grain products; mea, meat products; fat, fats and oils; sea, seasonings and condiments; leg, legumes, nuts, oilseeds and spices; fru, fruit products; veg, vegetable products; alc, alcoholic beverages; cof, coffee, cocoa and tea.

This frequency hierarchy was structured by chemical groups (**Figure S2**). Generalists were largely restricted to alcohols, aldehydes, and esters, whereas intermediaries spanned a broader range of structural classes. Individualists encompassed nearly all chemical groups, including several classes that were exclusively observed in this category, such as lactones, nitriles/amides, acetals, and halogens.

To examine how frequency-based roles differ within individual food categories, we profiled KFO frequency across 10 balanced, data-rich food categories (**Table S3**; see *Methods*), thereby minimizing biases arising from uneven category representation. The relative proportions of generalists, intermediaries, and individualists varied markedly across categories (Figure 2b). Meat products contained the largest number of category-level generalists, whereas plant-derived categories generally exhibited fewer generalists. These patterns parallel our earlier observations that plant categories exhibit higher diversity of KFO compositions, whereas animal-based groups show more constrained and similar chemical compositions.

Several food categories also displayed category-specific generalists (Figure 2d). Alcoholic beverages showed the highest proportion of exclusive generalists (50%), such as ethyl decanoate, ethyl 3-methybutanoate, and hexanoic acid, followed by seasonings and condiments (47%). In contrast, vegetables, legumes, nuts, spices, and fats and oils showed no exclusive generalists, with all generalists shared across multiple food categories.

Correlation analysis further clarified the functional roles of these frequency classes (Figure 2c). The number of generalists in a category was positively associated with within-category similarity (r=0.63, p=0.05), suggesting that these widely shared odorants form a backbone that stabilizes food aroma identity. Intermediaries showed a strong negative correlation (r=−0.66, p=0.04), indicating they drive aromatic divergence, whereas individualists showed no significant association (r=−0.06, p=0.88), consistent with their role in encoding product-specific nuance.

### Cross-category similarity in aroma composition reveals convergent patterns

To map compositional similarity among individual food aroma formulations in the KFO-Atlas, we projected them into a two-dimensional space using TMAP^17^, a tree-based dimensionality reduction method, in which each node represents an aroma formulation and spatial proximity reflects similarity in KFO composition (Figure 3; *see Methods*). The resulting map showed pronounced differences in spatial distribution across food categories. Alcoholic beverages formed a densely clustered region, whereas legumes, nuts, oilseeds, and spices occupied a highly dispersed region in the space.

**Figure 3.**
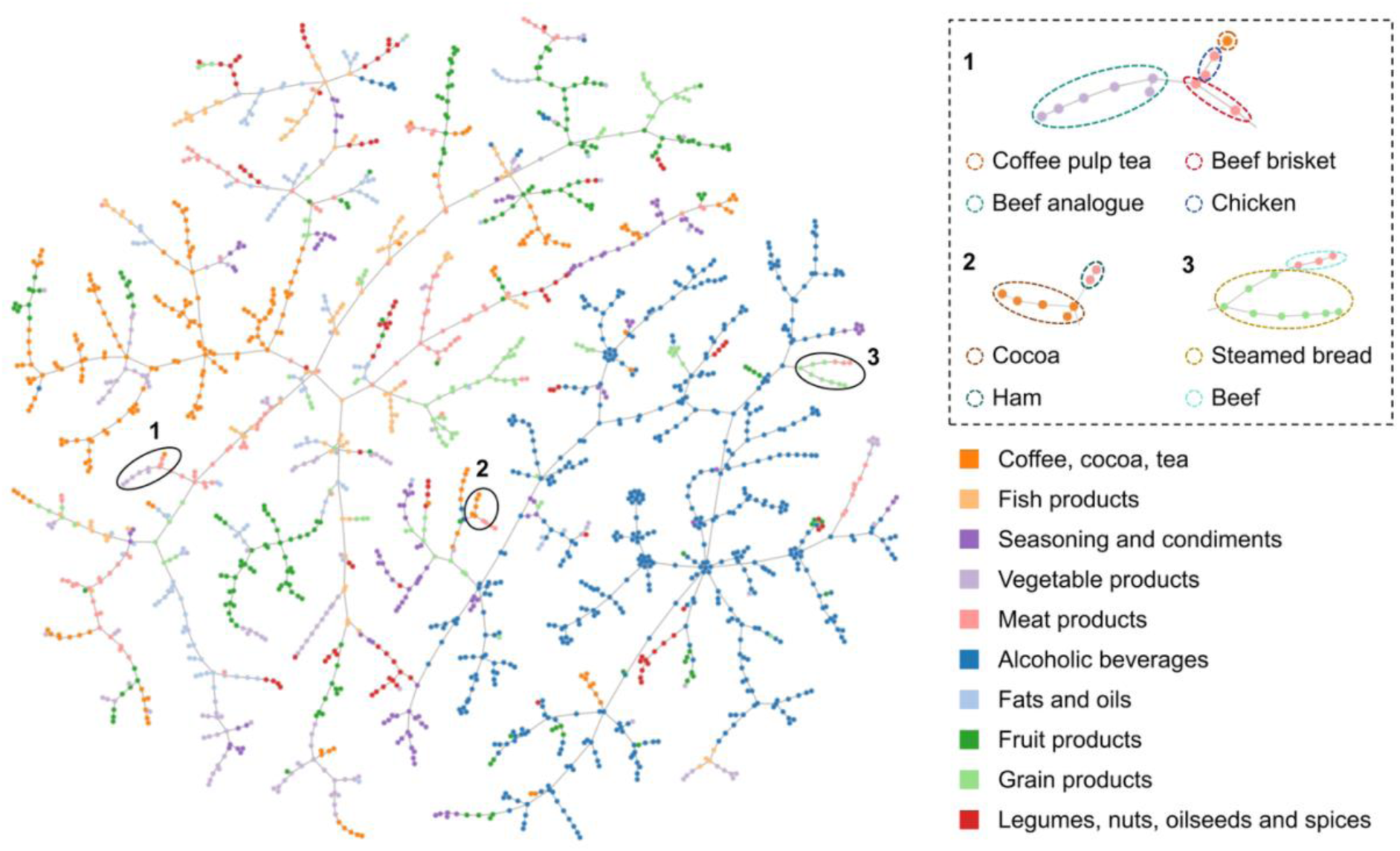
Cross-category neighborhoods in the KFO-Atlas aroma space. Aroma space of 1,876 aroma formulations based on key food odorants (KFO) composition, in which each node represents a formulation and spatial proximity reflects similarity in KFO profiles. Different food categories exhibit distinct spatial organizations: for example, alcoholic beverages form a densely clustered region, whereas legumes, nuts, oilseeds, and spices occupy a highly dispersed region of the map. Despite overall category separation, several annotated cross-category neighborhoods are observed. In the plot, we annotated Groups 1, 2, and 3. Group 1 includes formulations of plant-based meat analogues (light violet dots) and coffee pulp tea (orange dots) positioned adjacent to those of beef brisket and roasted chicken (pink dots). Group 2 shows selected cocoa products located near ham formulations. Group 3 places fermented Chinese steamed bread in close proximity to beef formulations.

Despite overall category-level separation, multiple cross-category neighborhoods were observed in the aroma space (Figure 3). In Group 1, a coffee pulp tea and plant-based meat analogues were positioned adjacent to beef brisket and roasted chicken because they shared several lipid-oxidation-derived odorants, including 1-hexanal, 2-pentylfuran, (E)-2-octenal, 1-octen-3-ol, and phenylacetaldehyde^18–21^. In Group 2, special cocoa products and ham clustered together due to shared Maillard- or lipid-oxidation odorants such as 3-methylbutanal, benzaldehyde, 2-heptanone, and 2-nonanone^22,23^.

Group 3 revealed proximity between beef and a fermented Chinese steamed bread. These formulations shared multiple odorants, including butanoic acid, styrene, 3-hydroxy-2-butanone, and benzothiazole^24–26^. The presence of butanoic acid, which is commonly associated with animal fats, within sourdough systems is potentially due to metabolic cross-talk between lactic acid bacteria and yeasts during fermentation, demonstrating the capacity of cereal-based fermentations to generate “meaty” olfactory notes.

Taken together, the analysis identifies instances in which formulations from different food categories occupy adjacent regions of the aroma space and share overlapping sets of KFOs. This convergence offers an explanation for why foods with different biological origins or culinary treatments may nonetheless exhibit related aromatic signatures. It also shows that plant-derived matrices, particularly those undergoing thermal or microbial transformation, may recreate meat-like aromas. This suggests the possibility of mimicking the aroma signatures of animal meat by assembling odorants found in plant-derived foods.

### Generative modeling enables targeted design of new aroma formulations

To explore whether aroma formulations can be computationally generated, we developed a generative framework based on a conditional variational autoencoder (CVAE) trained to learn the latent structure of KFO combinations and their relative proportions (Figure 4a). Guided by a multilayer perceptron classifier, the model generates formulations targeted to predefined aroma categories. The CVAE demonstrated stable convergence (validation loss=0.07; **Figure S3 and Table S4**), and the integrated multilayer perceptron classifier for post-hoc control achieved high predictive accuracy (ROC-AUC=0.96, mean average precision=0.85; **Figure S4**).

**Figure 4.**
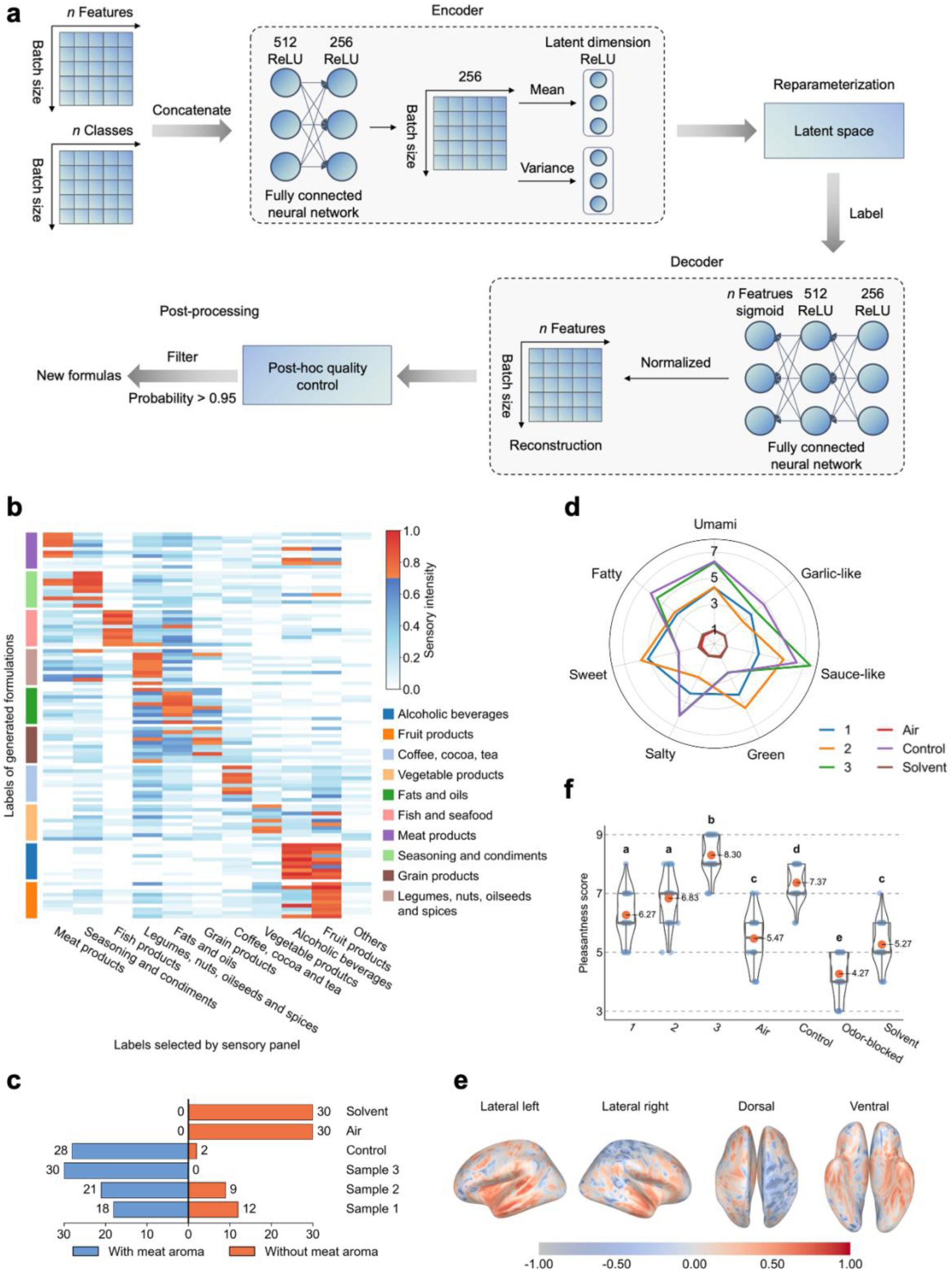
Generative reconstruction and multisensory evaluation of food aroma formulations. (a) Schematic of the conditional variational autoencoder used to generate KFO formulations targeting predefined aroma categories, with outputs further filtered by a multilayer perceptron classifier for category specificity. (b) Sensory evaluation of AI-generated formulations, showing the proportion of samples assigned by trained assessors to their intended aroma categories. (c) Sensory classification results for three AI-generated, plant-derived formulations targeting “meat”, with the proportion of assessors identifying each formulation as meat-like. (d) Sensory attribute profiles of sample 3 and the meat control across multiple aroma descriptors. (e) Electroencephalography source localization analysis comparing neural activation patterns elicited by sample 3 and the meat control, highlighting overlapping activated regions. Colors indicate the correlation scores. (f) Pleasantness ratings (1-9) obtained in an orthonasal-blocked tasting experiment using plant-based sausages, comparing samples with retronasal aroma enabled. Note: In panel (f), blue and orange scatter points represent individual pleasantness ratings assigned by participants. Gray box plots summarize the data distribution for each sample, indicating the first and third quartiles, the median, and the range of non-outlying values. Dark red points denote the mean pleasantness for each sample, and black curves represent the corresponding probability density distributions.

When generating 1,000 candidate formulations per category under a 5-20 KFO constraint, the framework produced 853 high-confidence outputs (classifier probability>0.95). The average maximum similarity between generated and existing formulations within the target category was 0.58, indicating the ability to generate novel combinations (**Table S5**). To assess perceptual validity, 30 experienced panelists evaluated 100 randomly selected AI-generated formulations (10 per food category) in a blinded setting (**Supporting Data 2**). Across categories, 73% of generated formulations were assigned by panelists to their intended aroma category (Figure 4b).

Inspired by the cross-category convergence observed earlier, we next tested whether the model could recreate meat-like aromas under a source-constrained design scenario, in which only KFOs previously reported in plant-derived food formulations were allowed. Among 730 generated candidates, three formulations composed exclusively of plant-derived KFOs were randomly selected for sensory evaluation (**Table S6**). Two formulations were classified as meat-like by 65-70% of assessors, whereas one formulation (sample 3) was classified as meat-like by all assessors (Figure 4c). Duplicate evaluations showed no significant differences between repeated measurements (paired-sample t-tests; **Figure S5**). Sensory profiling further showed that sample 3 exhibited attribute intensities similar to the aroma profile of meat control from literature^27^ across multiple descriptors (Figure 4d), while relying on a reduced number of KFOs derived from plant-based sources (**Table S6**).

To further compare neural responses elicited by sample 3 and the meat control^27^, EEG source localization analysis was performed (Figure 4e). Overlapping activation patterns were observed across multiple brain regions, including the parahippocampal gyrus (r=0.46), insula (r=0.45), superior temporal gyrus (r=0.43), and posterior cingulate cortex (r=0.43), indicating similar neural response profiles between the two stimuli. Such correlation magnitudes are commonly interpreted as meaningful neural similarity in human electrophysiological and neuroimaging studies^28^.

To evaluate the contribution of the designed aroma formulation to pleasantness, an orthonasal-blocked tasting experiment was conducted using plant-based sausages. Allowing retronasal aroma resulted in higher pleasantness scores compared to aroma-blocked conditions, showing the importance of aroma enhancement for plant-based meat alternatives (Figure 4f). Among the evaluated aromas, sample 3 had the highest pleasantness score and reached values above those reported for the real meat control^27^.

## Discussion

In this study, we constructed KFO-Atlas, a comprehensive resource comprising 896 key food odorants spanning 20 food categories and 87 subcategories. By substantially expanding previous repertoires of KFOs, KFO-Atlas moves beyond earlier estimates that only approximately 230 food volatiles account for most odor-active components. This expanded atlas provides a more robust foundation for examining how food aromas are organized, compared, and related across diverse foods.

Analysis of KFO-Atlas revealed a structured hierarchy in which generalists, intermediaries, and individualists play complementary functional roles. Generalists form a shared backbone that stabilizes food aroma identity, intermediaries drive divergence, and individualists capture product-specific nuances. Together, this hierarchy indicates that food aroma is structured and interpretable rather than arbitrary.

These organizational rules manifest differently across food categories. Plant-derived foods, shaped by diverse secondary metabolism, exhibit highly heterogeneous odorant compositions. In contrast, animal-derived and fat-rich foods repeatedly converge on lipid-oxidation- and Maillard-derived odorants, resulting in more constrained and similar compositions. Importantly, cross-category neighborhoods in the KFO-Atlas show that distinct foods can, in specific cases, assemble similar odorant combinations through shared reaction pathways. This convergence provides a mechanistic explanation for long-recognized sensory similarities among different foods.

This structural convergence has direct implications for aroma design. Food categories governed by constrained chemical compositions occupy relatively limited regions of the KFO space, rendering their aroma profiles more learnable and reconstructable. Meat products exemplify this pattern: despite diversity in raw materials and processing conditions, their aromas consistently converge on a limited set of odorants.

Building on this structural property, we trained a conditional variational autoencoder to learn the latent organization of KFO combinations and generate category-targeted aroma formulations. Blinded sensory evaluation showed that most AI-generated formulations were perceptually recognized as their intended categories, indicating that the model captures perceptually meaningful organization rather than compositional similarity. This advances beyond conventional AI approaches focused on individual odorants, addressing a longstanding challenge in modeling the emergent sensory properties of multi-odorant mixtures. In this respect, our results complement recent efforts in odorant perception prediction and mixture modeling^12,29^ by demonstrating that perceptually coherent mixture organization can be learned and validated at the level of real-world odorant combinations.

The relative tractability of meat aroma provided a meaningful test case for generative design, as meat products combine strong perceptual expectations with convergent KFO compositions. Using exclusively plant-derived KFOs, the model generated formulations that were consistently perceived as meat-like by human assessors. Notably, multisensory validation showed that one plant-derived formulation elicited neural activation patterns overlapping with those induced by authentic meat aroma across canonical olfactory and affective brain regions. This convergence at the neural level suggests that shared odorant architectures can engage higher-level perceptual processing in a manner comparable to animal-derived aromas, rather than merely approximating sensory cues. Beyond categorical recognition, aroma reconstruction exerted a measurable influence on consumer-relevant perception. Allowing retronasal aroma substantially increased pleasantness ratings of plant-based sausages, and the best-performing plant-derived aroma formulation exceeded the pleasantness of the real meat control. Together, these findings indicate that generative aroma design can not only reproduce established sensory profiles but also access regions of aroma space that may remain underexplored in conventional formulation strategies.

Several limitations point to directions for future development. Although KFO-Atlas substantially expands the reported landscape of key food odorants, it remains shaped by biases in the literature, including variability in analytical protocols, sample preparation, and population-specific odor thresholds. These factors introduce noise into KFO identification and odor activity value estimation and may affect different food categories unevenly. The generative model is likewise constrained by data availability, with categories represented by fewer formulations yielding less stable latent representations and, consequently, less robust outputs. In addition, the FoodEx2^16^ classification system, while comprehensive, was not designed for aroma analysis and contains overlaps that may obscure finer sensory distinctions. Future efforts would benefit from aroma-oriented food ontologies, expanded and more balanced formulation datasets, and the integration of practical constraints, such as ingredient availability, cost, and manufacturing feasibility, to bridge the gap between conceptual generative design and deployable aroma formulation.

KFO-Atlas and the generative framework together provide a set of complementary resources with broad relevance across food science and innovation. At a fundamental level, they offer a mechanistic understanding of how odorants interact to shape aroma identity, diversity, and convergence. At an applied level, off-notes are a major obstacle to the development of alternative protein sources^30,31^. This framework addresses this problem by enabling more rational aroma formulation, accelerated prototyping, and the development of alternative protein products that capture key sensory attributes traditionally associated with animal-derived foods. Beyond formulation, a structured view of odorant prevalence and functional roles may also support greater transparency in ingredient assessment and flavor labelling.

Together, KFO-Atlas and the generative modeling framework establish a data-driven foundation for decoding and designing food aroma. By revealing when and why aroma becomes perceptually reconstructable at the level of key odorant combinations, this work enables a more systematic and predictive approach to flavor science and food innovation.

## Methods

### Data collection and curation

KFOs were defined as volatile compounds with odor activity values (OAVs) greater than one^6^. To assemble a comprehensive and up-to-date dataset of KFOs, we conducted a structured literature survey of food aroma studies published between January 2015 and July 2024. Publications were retrieved from Google Scholar using a Boolean keyword strategy (“food” AND “aroma” AND “GC” AND “OAV”), yielding 5,892 candidate records. Titles and abstracts were initially screened using FoodBERT^32^, a domain-specific language model trained to identify food-related content, to remove clearly irrelevant studies reporting non-food systems or lacking aroma analysis. All remaining records were subsequently examined by manual inspection. After this two-stage filtering, 1,667 publications reporting specific food items or ingredients and quantitative aroma analysis were retained for data extraction.

For each eligible study, KFOs were manually curated and cross-validated. Extracted information included food identity, lists of KFOs detected under the reported experimental conditions, corresponding OAVs, odor thresholds, and quantitative concentrations when available. To ensure consistency across studies, all compounds were standardized using the PubChem ID identifier. As OAVs reported in the literature vary substantially due to differences in food samples and experimental conditions, we retained the OAVs as reported in the original studies, without attempting cross-study re-standardization. Accordingly, KFOs were consistently identified based on a relative criterion (OAV > 1) within each study, reflecting compounds perceived as aroma-active under their specific food sample and experimental context.

Each aroma formulation was assigned to a food category using the FoodEx2^16^ classification system, resulting in 20 major food categories and 87 subcategories. This hierarchical annotation enabled consistent comparison across diverse food systems. For example, meat products were further resolved into mammal meat, bird meat, dried meat, cooked cured meat, raw cured meat, and preserved sausages. Food items that could not be unambiguously mapped to an existing subcategory (for example, composite fermented cereal-dairy products such as Tarhana) were assigned at the major-category level only to avoid introducing artificial classification bias.

To characterize how different odorants contribute to food aroma organization, KFOs were classified according to their occurrence frequency across formulations, following and extending a previously established framework^7^. Compounds detected in more than 25% of formulations were designated generalists, those occurring in 5-25% were defined as intermediaries, and those present in fewer than 5% of formulations were classified as individualists.

All KFOs were additionally assigned to one of 18 chemical classes based on molecular structure, using the Volatile Compounds in Food reference classification (https://www.vcf-online.nl/VcfCompounds.cfm). These classes included alcohols, aldehydes, ketones, esters, acids, furans, phenols, sulfur compounds, hydrocarbons, bases, lactones, (ep)oxides/pyrans/coumarins, ethers, acetals, nitriles/amides, oxazol(in)es, halogens, and imidazoles (**Table S1**).

### Clustering of food aroma formulations

To characterize the organizational structure of aroma compositions across foods, we restricted hierarchical analyses to food categories represented by at least ten aroma formulations, ensuring sufficient sampling to yield stable similarity estimates. This filtering resulted in a curated dataset of 2,274 formulations for comparative analysis. Compositional relationships among formulations were visualized using hierarchical clustering implemented with PyComplexHeatmap^33^. Clustering was performed on proportional KFO composition vectors using correlation distance and average linkage. Correlation distance was selected to emphasize similarities in relative odorant patterns rather than absolute KFO abundance, allowing formulations with comparable compositional structure to cluster together despite differences in scale.

### Similarity calculation between food aroma formulations

To quantify compositional similarity between aroma formulations, we designed a weighted similarity metric that jointly captures similarity in odorant composition patterns and differences in relative abundance. Specifically, we integrated cosine similarity, which reflects directional similarity in KFO composition, with Manhattan distance, which captures absolute proportional deviations between formulations. For two formulations *A_i_* and *B_i_*, all KFOs identified across the dataset were first expanded into a unified feature vector, with absent compounds assigned a value of zero. Each vector element represents the proportional contribution of a given KFO within the formulation. Similarity between formulations was then computed as:

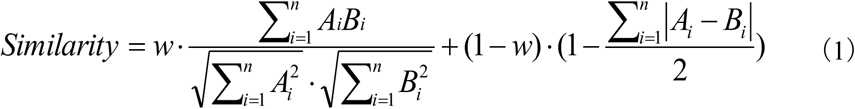

where the weighting coefficient *w* = 0.5 assigns equal weight to compositional pattern similarity (cosine term) and proportional deviation (Manhattan term). The resulting similarity scores range from 0 to 1, with higher values indicating greater overlap in KFO composition.

For each food category, within-category similarity was defined as the mean of all pairwise similarities among formulations belonging to that category. This provides a robust estimate of category-level aromatic homogeneity while accounting for differences in both KFO identity and relative abundance.

### Hierarchical roles of key food odorants in shaping food aroma

To resolve category-specific frequency patterns while ensuring statistical robustness, we restricted the analysis to food categories represented by at least 50 aroma formulations, and to subcategories with at least 20 formulations. This filtering yielded a balanced, data-rich subset comprising ten well-represented food categories (**Table S3**). Formulations corresponding to composite dishes were excluded because they integrate multiple ingredient types and confound category-level interpretation. In addition, fruit and vegetable juices and nectars were removed owing to their substantial raw-material overlap with whole fruit and vegetable categories in the FoodEx2^16^ classification.

Within this curated dataset, KFOs were classified as generalists, intermediaries, or individualists based on their occurrence frequency, and the abundance of each class was quantified at the category level. To assess how different KFO frequency classes relate to category-level aroma organization, we computed Spearman’s rank correlations between the number of generalist, intermediary, and individualist KFOs in each category and the corresponding within-category similarity scores. Spearman correlation was chosen to account for non-normal distributions and the limited number of categories analyzed. This analysis quantifies the extent to which different classes of KFOs contribute to aromatic convergence or heterogeneity across food categories.

### Cross-category convergence analysis

To examine cross-category convergence among aroma formulations, we constructed a global aroma space across the ten balanced, data-rich food categories (**Table S3**) using TMAP^17^, a tree-based dimensionality-reduction method optimized for sparse data. TMAP preserves local neighborhood relationships, making it well-suited for identifying convergent molecular architectures among complex odorant mixtures.

For each formulation, the proportional contributions of all KFOs were scaled to a uniform range and discretized into 8-bit unsigned integer vectors, satisfying the input requirements of the TMAP hashing pipeline while preserving relative compositional information. The resulting vectors were embedded in a tree-structured map that encompassed all eligible formulations. Nodes were colored by food category, and representative products were annotated to facilitate interpretation. In this global aroma map, tightly packed branches reflect strong compositional convergence within food categories. In contrast, mixed branches indicate cross-category molecular overlap, highlighting instances where distinct food matrices converge on similar odorant architectures.

### Development of supervised machine learning models

To enable post hoc quality control of generated aroma formulations, we developed supervised classification models to predict food category membership based on KFO composition. Using the ten balanced, data-rich food categories, together with an additional ‘others’ class representing chemically plausible but category-agnostic formulations (*see Supporting Information*), we benchmarked multiple classifiers, including linear discriminant analysis, decision trees, support vector machines, logistic regression, and multilayer perceptron. Formulations were represented by their proportional KFO composition vectors and labeled according to food categories, with the ‘others’ class introduced to capture off-category formulations and improve robustness. Among the tested approaches, the multilayer perceptron achieved the best overall performance (macro F1=0.87 ± 0.02; **Table S7**) and was therefore selected as a probabilistic filter to screen CVAE-generated formulations, ensuring category fidelity in downstream analyses. Full benchmarking details are provided in the *Supporting Information*.

### Development of a conditional variational autoencoder

To enable de novo generation of aroma formulations targeted to specific food categories, we developed a generative framework based on a CVAE, integrated with a supervised classifier for post hoc quality control. The model was trained on formulations from ten balanced, data-rich food categories, together with an additional ‘others’ class representing chemically plausible but category-agnostic formulations. Input features consisted of proportional KFO composition vectors. The CVAE encoder jointly processes the KFO composition vector and the corresponding category label to learn a latent representation that captures category-conditioned structure in KFO combinations. The encoder outputs the mean (μ) and log-variance (log σ²) parameters of a multivariate Gaussian latent distribution, from which latent variables are sampled using the reparameterization trick. The decoder receives the sampled latent vector and the category label, and reconstructs a formulation vector using multilayer perceptrons with ReLU activations, followed by a sigmoid output layer that constrains all values to the [0,1] interval.

Because empirical aroma formulations are sparse and dominated by a limited number of key odorants, we applied a sparsity-preserving post-processing step to all generated outputs. For each decoded formulation, only the top k KFOs (with k constrained between 5 and 20 during validation) were retained, and the resulting vectors were renormalized to sum to one. This procedure yields reasonable formulations that reflect the structure observed in real foods.

To ensure that generated formulations matched the intended aroma category, we applied a previously trained multilayer perceptron classifier as a probabilistic filter. Only formulations assigned to the target category with high confidence (probability>0.95) were retained. This post hoc filtering step does not participate in generation but serves to remove off-category outputs and improve perceptual consistency in downstream evaluation.

The CVAE was trained using a composite objective comprising a reconstruction term and a Kullback-Leibler divergence regularization term. Reconstruction loss was computed using mean squared error between the input and reconstructed formulation vectors (**eq. 2**), whereas the Kullback-Leibler term encouraged the latent space to approximate a standard normal prior **(eq.3)**. The total loss **(eq.4)** was represented as:

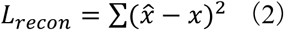

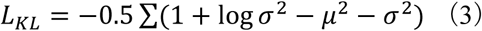

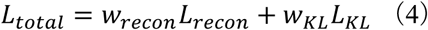

The AdamW optimizer was used for parameter updates. A cosine annealing schedule was used to modulate β across epochs, while a ReduceLROnPlateau scheduler halved the learning rate when the validation loss plateaued (patience=20 epochs). Gradient clipping (maximum norm=1.0) stabilized training, and early stopping was triggered when validation loss failed to improve over the specified patience epochs. To identify an optimal architecture and training configuration, we conducted an extensive hyperparameter search (1,000 trials), with each trial capped at 1,000 epochs. The best-performing model was selected based on minimum validation loss. Full hyperparameter ranges are summarized in **Table S4**.

To quantify the novelty of generated formulations relative to existing data, we calculated a maximum-average similarity metric. For each generated formulation, its highest similarity to any formulation within the target category was identified and averaged across samples. Lower similarity scores indicate exploration of novel yet category-consistent regions of the aroma formulation space.

### Sensory and electroencephalography assessment

Thirty experienced panelists were recruited for sensory evaluation. All participants were right-handed, reported no respiratory, neurological, or olfactory disorders, and had no history of substance abuse. The study was conducted in accordance with institutional ethical guidelines and was approved by the Ethics Committee of the Shanghai Institute of Technology (approval number: SIT-2024-LL26).

Sensory evaluation comprised two sections. In Section 1, a blinded category-recognition task was used to assess whether the generative model could produce aroma formulations that were perceptually consistent with their intended food categories. In Section 2, a focused evaluation examined whether meat-like aroma formulations reconstructed exclusively from plant-derived KFOs could reproduce characteristic meat aroma profiles and influence hedonic perception.

In Section 2, a focused tasting experiment was conducted to evaluate the contribution of aroma to pleasantness perception during consumption. Pleasantness scores range from 1 to 9, with 9 indicating the highest level. Selected meat-targeted aroma formulations composed exclusively of plant-derived KFOs were incorporated into a standardized plant-based sausage matrix, and pleasantness was assessed under aroma-available and aroma-blocked conditions.

To further probe perceptual similarity at the neural level, EEG recordings (**Figure S6**) were obtained from a subset of participants during aroma exposure to selected plant-derived and animal-meat formulations. Source localization and correlation analyses were used to compare neural activation patterns associated with aroma perception. Full experimental procedures and data processing pipelines for sensory and EEG analyses are provided in the *Supporting Information*.

## Acknowledgements

This project was supported by the Collaborative Innovation Center of Fragrance Flavour and Cosmetics, Ministry of Education, Singapore, under the Academic Research Fund Tier 1 (A-8003718-00-00), and NUS IT (NUSREC-HPC-00001). The authors acknowledged using an AI tool (ChatGPT) for grammar check and are fully responsible for the content and conclusions of the manuscript.

## Author contributions

D.Z. X.K. and Q.K. designed the research. J.Z. and D.Z. collected and analyzed the data. J.Z. and H. X. developed and evaluated the machine learning models. J.Z. implemented the experiments. D.Z. and J.Z. wrote the paper with input from A.P., X.K. and Q.K. All authors approved the final paper.

## Data availability

Data generated in this research are available in Supporting Data 1-2 and Supporting Information. Full data from the KFO-Atlas will be publicly available in a Zenodo repository (https://zenodo.org/records/18311787).

## Code availability

Code for the CVAE model reported in this research is available in a Zenodo repository (https://zenodo.org/records/18333737).

