## Supplemental methods, figures, and tables for "Molecular atlas of key food odorants reveals structured aroma organization and enables generative aroma design"

* Correspondence:

### Supplementary methods

#### Development of supervised machine learning models

Formulation vectors, including randomly generated samples assigned to the 'others' class, were encoded as proportional key food odorants (KFOs) composition spanning all identified KFOs. Five supervised learning algorithms, decision trees, support vector machines, logistic regression, linear discriminant analysis, and multilayer perceptron, were evaluated for classification performance.

Models were trained five times using five independent random seeds (100, 200, 300, 400, and 500). Performance was assessed using weighted F1 scores and area under the ROC curve (ROC-AUC). Across all evaluated settings, the multilayer perceptron achieved the highest performance (AUC>0.90 across categories; F1 score=0.87; Figure S4; Table S7). This model was therefore selected for filtering CVAE-generated formulations.

#### Generation of formulations in the 'others' category

To ensure the classification and generation models are not forced to assign one out of the ten groups to a formulation, we created a 'others' category in the training dataset. For each formulation, a random integer between 3 and 91 (Figure 1a), corresponding to the observed minimum and maximum numbers of KFOs per formulation, was selected. The corresponding number of KFOs was randomly sampled, assigned weights at random, and normalized to sum to 1. Each generated formulation was represented as a sparse vector over all KFOs. To avoid potential overlaps and ensure divergence from existing formulations, the average similarity to all empirical formulations was computed, and only formulations with an average similarity below 0.2 were retained. Using this procedure, 500 formulations were generated for model training.

#### Sensory evaluation

In Section 1, to evaluate whether AI-generated aroma formulations were perceptually recognizable, 100 formulations were selected from model outputs across ten food categories. Within each category, ten formulations were randomly sampled using a fixed random seed (42) to ensure reproducibility (Supporting Data 3). All samples were prepared according to their generated recipes and diluted to 5% (w/w) in 1,2-propanediol. Samples were presented in identical containers coded with three-digit blind identifiers.

Testing was conducted in a well-ventilated laboratory. Before the experiment, participants were fully informed of the procedures and confirmed their voluntary participation. During evaluation, each participant was presented with a scented strip corresponding to a coded sample and instructed to select the matching food categories from a set of 11 predefined options (10 food categories plus ‘others’). For each sample, participants could select one to three categories, marking the selected categories as 1 and the unselected categories as 0.

In Section 2, to assess whether meat-like aromas could be reconstructed using exclusively plant-derived KFOs (e.g., those present in coffee, cocoa, tea, fruits, grain products, legumes, nuts, oilseeds, spices, and vegetables), three formulations were randomly selected from 730 generated “meat” candidates composed only of plant-origin KFOs and designated as Samples 1-3. Sample 3 was duplicated to assess panel consistency. All formulations were prepared according to their recipes (Table S6) and diluted to 5% (w/w) in 1,2-propanediol. All compounds were sourced from Shanghai Meixin Chemical Technology Co., Ltd. A blank sample (air) and a matrix sample (1,2-propanediol) were also included, resulting in a total of seven samples for evaluation. Participants rated each sample across eight dimensions of the aroma wheel, following the design of Wang et al.[1], including umami, garlic-like, sauce-like, green, salty, sweet, meaty, and fatty, using a 1-9 scale.

For hedonic evaluation, to minimize the influence of movement on flavor perception, aromas were delivered to participants’ faces at a flow rate of 5 L/min using a gas pump positioned 5-7 cm from the nose. The experimenter first stabilized aroma delivery with the pump, then provided a standardized plant-based ham sausage sample, instructing participants to chew and swallow. Immediately after swallowing, participants rated hedonic response using the SAM scale[2]. To investigate the contribution of orthonasal olfaction to hedonic perception, a control condition was included in which participants wore nasal clips to block orthonasal airflow while performing the same tasting and rating procedure. To prevent cumulative effects of taste and smell, a 5-minute interval was set between trials, during which participants were given small amounts of warm water and briefly removed from the testing area to clear the oral and nasal passages. Plant-based ham sausages were purchased from Shenzhen Whole Perfect Foods Co., Ltd., contained no added aroma, and were uniformly cut into 1.5 cm pieces.

Panelists were not formally trained prior to the sensory evaluation. This choice was motivated by practical considerations. Unlike well-defined aromas such as “apple” or “vanillin”, the target stimuli in this study correspond to broad and chemically heterogeneous food categories (e.g., fruits, vegetables, beverages, and meat), for which no standardized reference compounds or canonical training samples exist. In addition, the aroma identification (e.g., meat-like versus non-meat-like) focuses on category-level recognition and does not require fine-grained, calibrated descriptor scaling or expert cue learning. These tasks rely on holistic perceptual assessment rather than analytical discrimination. Therefore, we used experienced (having participated in at least 3 sensory evaluations) but untrained panelists to provide a conservative test of whether the learned KFO organizations translate into perceptually recognizable aroma identities under natural smelling conditions.

**Electroencephalography assessment**

Participants were seated comfortably in ergonomically designed chairs. Aromas of sample 3 and the control were delivered using a gas pump at a flow rate of 5 L/min, positioned 5-7 cm in front of the participant's face (Figure S5a). EEG recordings consisted of a 30-second resting period followed by a 60-second aroma-exposure period (Figure S5b). Participants were instructed to press the green button on the SAGA synchronization module immediately upon perceiving the aroma, marking the onset of the exposure period.

EEG data were acquired using a TMSI SAGA EEG device (Twente Medical Systems International BV, the Netherlands) with a 64-channel cap arranged according to the international 10-20 system (Figure S5c) and sampled at 500 Hz. Electrode impedance was maintained below five kΩ.

Preprocessing was performed in MATLAB using the EEGLAB toolbox [3]. Signals were band-pass filtered between 1 and 70 Hz, and a 50 Hz notch filter was applied to remove power-line interference. Independent component analysis was used to remove artifacts, including electrooculographic and electromyographic activity. Channels M1 and M2 were excluded, and an average reference was applied. Based on the recorded event times, the subsequent 30 s of data were extracted as aroma exposure segments, which were further divided into 2 s epochs with 50% overlap [4].

All subsequent analyses were performed using the MNE-Python software package [5]. To ensure accurate visualization and source modeling, the standard fsaverage brain template was used, providing a normalized cortical surface and source space for group-level analysis [6].

For source localization, forward models and noise covariance matrices were computed separately for each experimental condition. Forward models were generated using a boundary element model based on the fsaverage template, with the source space defined on the cortical surface of fsaverage, containing 10,242 vertices per hemisphere at ico-5 resolution. Forward models combined EEG sensor locations, boundary element model, and source space, with a minimum source spacing of 5 mm to avoid closely spaced sources. Noise covariance matrices were estimated primarily from the full set of epochs, with regularization parameters optimized automatically. Source localization was performed using dynamic statistical parametric mapping.

For each condition, EEG epoch data were loaded from preprocessed files and aligned to the standard 10-20 electrode layout. The inverse operator was configured with loose orientation constraints (loose = 0.2) and depth weighting (depth = 0.8) to account for deeper cortical sources. A signal-to-noise ratio (SNR) of 3.0 was assumed to calculate the regularization parameter ($\lambda^2 = 1/SNR^2$). Source time courses (STCs) were computed for each epoch and subsequently averaged across epochs within each condition to produce mean STCs for statistical analysis.

To examine differences in brain activity across conditions, the mean STCs were compared using correlation analyses. For each vertex in the source space, time series from two conditions were compared using Pearson or Spearman correlation, depending on data normality, assessed with the Shapiro-Wilk test. Given the exploratory nature of this analysis, vertex-wise significance was used to define candidate regions, and results are interpreted at the level of regional patterns rather than single-vertex inference. Vertices with $p < 0.05$ were identified as significant. Significant vertices were mapped to anatomical labels using the aparc parcellation of the fsaverage template, and for each label, the mean correlation coefficient and number of significant vertices were calculated.

For visualization, source-space similarity maps were created in MNE-Python using correlation coefficients as values. Maps were rendered on the fsaverage cortical surface with a symmetrical color scale from -1 to 1. Four 3D views (left lateral, right lateral, dorsal, and ventral) were generated using the PyVista backend[7].

**Table S1. Number and proportion of KFOs classified by chemical groups.**

| Chemical groups | Freqs | Proportions |
|---|---|---|
| Esters | 179 | 20.0% |
| Alcohols | 121 | 13.5% |
| Carbonyls, aldehydes | 99 | 11.0% |
| Carbonyls, ketones | 98 | 10.9% |
| Sulfur compounds | 91 | 10.2% |
| Hydrocarbons | 77 | 8.6% |
| Bases | 58 | 6.5% |
| Furans | 39 | 4.4% |
| Acids | 34 | 3.8% |
| Phenols | 31 | 3.5% |
| (Ep)oxides, pyrans, coumarins | 27 | 3.0% |
| Lactones | 23 | 2.6% |
| Nitriles and amides | 9 | 1.0% |
| Ethers | 4 | 0.4% |
| Acetals | 2 | 0.2% |
| Halogens | 2 | 0.2% |
| Imidazoles | 1 | 0.1% |
| Oxazol(in)es | 1 | 0.1% |

**Table S2. Number of KFOs per food aroma formulation across food categories.**

| Food categories | Mean | Max | Min |
|---|---|---|---|
| Alcoholic beverages | 13.38 | 57 | 3 |
| Coffee, cocoa and tea | 16.49 | 91 | 4 |
| Fruit products | 11.45 | 46 | 3 |
| Vegetable products | 13.71 | 37 | 4 |
| Fats and oils | 14.02 | 37 | 3 |
| Seasoning and condiments | 17.05 | 48 | 3 |
| Juices and nectars | 12.14 | 33 | 3 |
| Fish and seafood | 10.63 | 27 | 3 |
| Meat products | 14.88 | 33 | 3 |
| Composite dishes | 15.91 | 43 | 3 |
| Grain products | 11.71 | 45 | 3 |
| Legumes, nuts, oilseeds and spices | 17.02 | 47 | 4 |
| Food imitates and supplements | 13.12 | 34 | 3 |
| Milk and dairy products | 11.29 | 25 | 3 |
| Sugar and confectionery products | 13.27 | 24 | 3 |
| Major isolated ingredients and additives | 13.53 | 22 | 7 |
| Water-based beverages | 9.36 | 13 | 5 |

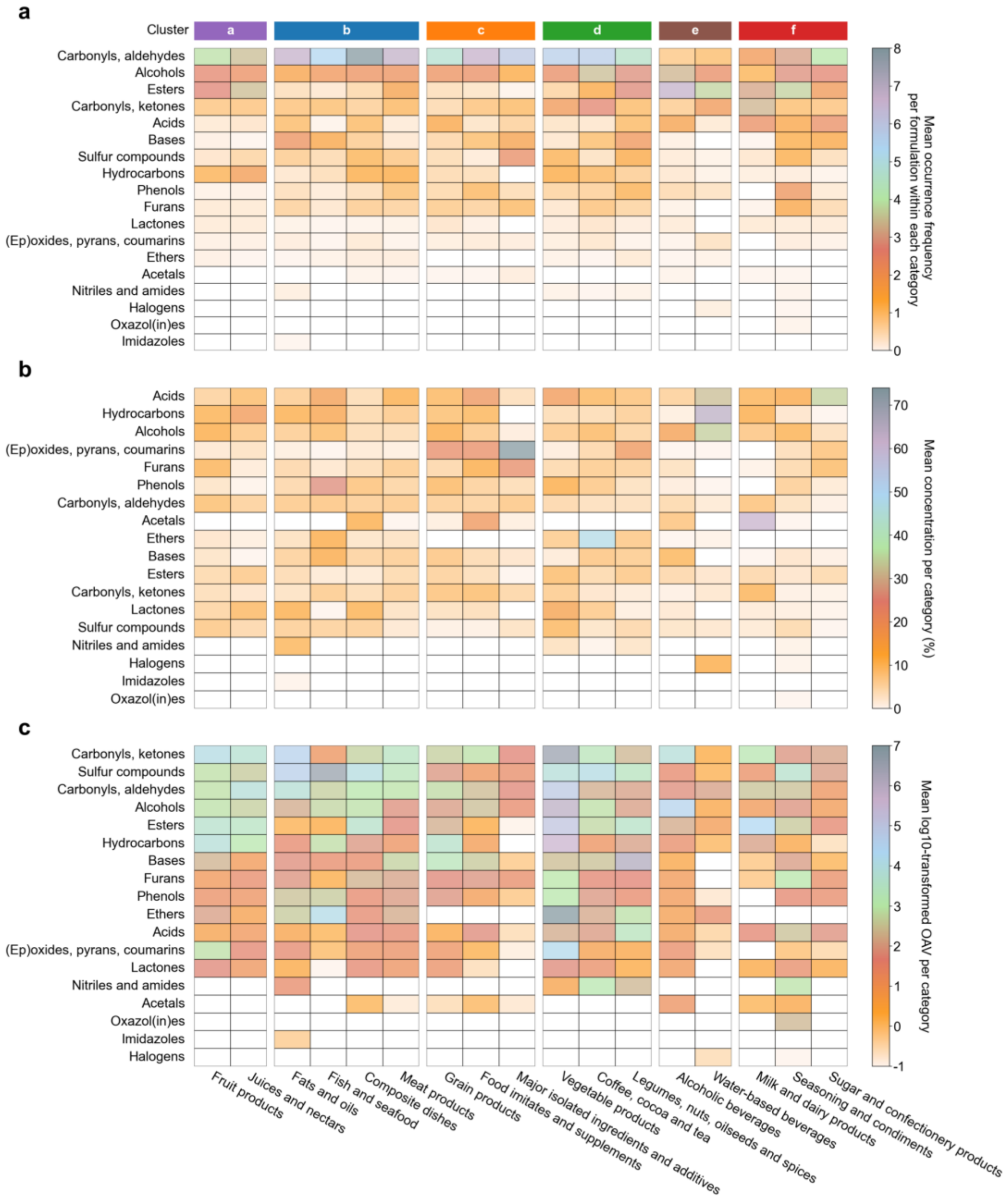


**Figure S1. Structural characterization of KFO compositions across food categories.** Heatmaps summarize the distribution of 18 chemical groups across 17 food categories, quantified by (a) mean occurrence frequency per KFO formulation, (b) mean relative concentration within each category, and (c) $\log_{10}$-transformed odor activity values (OAVs). Structural classes are ordered from top to bottom by their overall abundance across categories, facilitating comparisons of dominant and rare compound classes. White cells indicate the absence of the corresponding chemical groups in a given category.

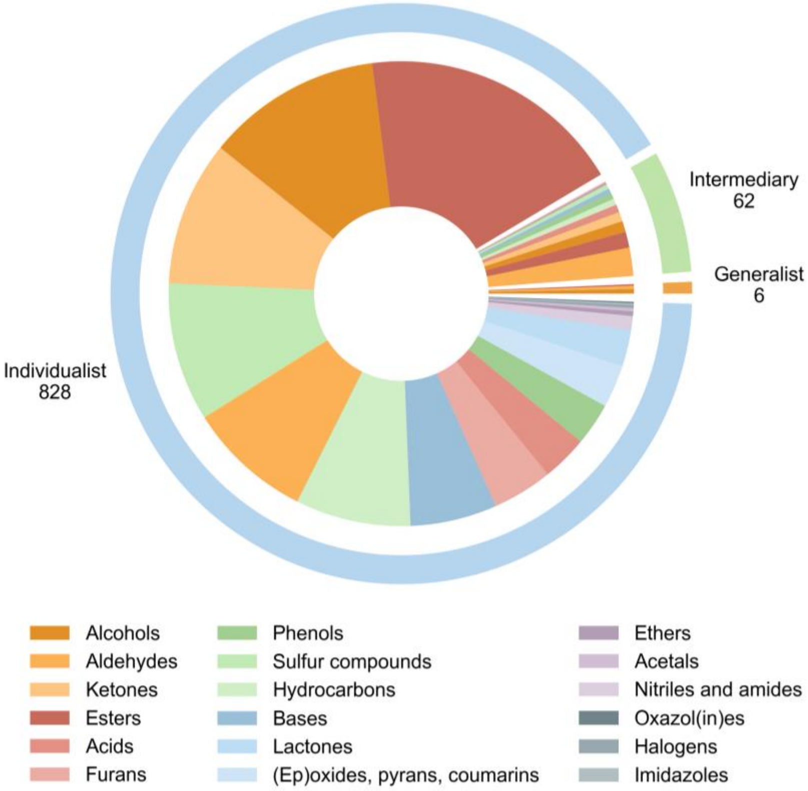


**Figure S2. Chemical class composition across frequency-defined KFOs groups.** Nested donut chart showing the chemical class composition of KFOs stratified by frequency across formulations. KFOs were classified into generalists (present in >25% of samples), intermediaries (5-25%), and individualists (<5%). The outer ring indicates the total number of KFOs in each frequency class, whereas the inner segments depict the relative contributions of different chemical classes. Generalists are confined to a small number of chemically simple classes, predominantly alcohols, aldehydes, and esters. Intermediaries span a broader range of structural classes, while individualists encompass all chemical classes.

**Table S3. Number of formulations for ten balanced, data-rich food categories.**

| Food categories | Counts |
|---|---|
| Alcoholic beverages | 546 |
| Fats and oils | 132 |
| Coffee, cocoa and tea | 236 |
| Fish and seafood | 125 |
| Fruit products | 200 |
| Grain products | 117 |
| Legumes, nuts, oilseeds and spices | 99 |
| Meat products | 125 |
| Seasoning and condiments | 130 |
| Vegetables products | 166 |
| Others | 500 |

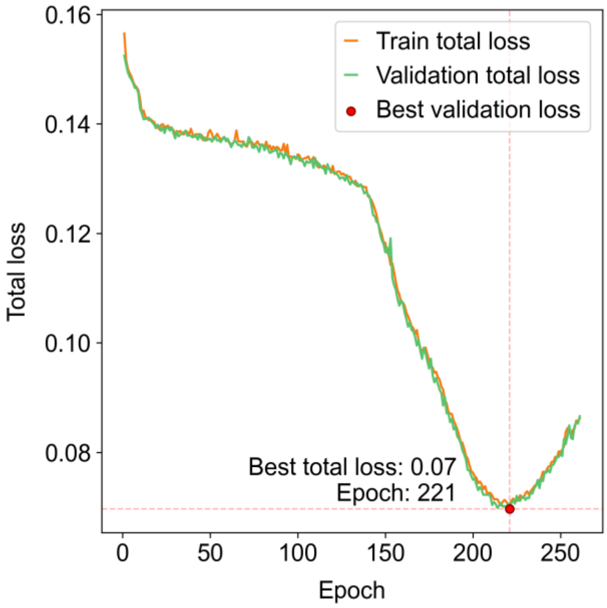


**Figure S3.** Training loss of the conditional variational autoencoder under the optimal hyperparameter configuration.

**Table S4. Hyperparameter search space for conditional variational autoencoder optimization.**

| Hyperparameters | Search ranges | Bests |
|---|---|---|
| latent_dim | [16, 32, 64, 128] | 64 |
| learning_rate | [1e-5, 5e-5, 1e-4, 5e-4] | 0.0005 |
| patience | [20, 30, 50] | 50 |
| recon_weight | [0.5, 1.0, 2.0] | 0.5 |
| kl_weight | [0.1, 0.5, 1.0] | 0.1 |
| sim_weight | [0.5, 1.0, 2.0] | 0.5 |
| beta_min | [0.05, 0.1, 0.2] | 0.05 |
| beta_max | [0.5, 1.0, 2.0] | 0.5 |
| beta_cycle | [50, 100, 200] | 200 |
| encoder_hidden | [512, 256]; [256, 128]; [512, 256, 128] | [512, 256, 128] |
| decoder_hidden | [256, 512]; [128, 256]; [128, 256, 512] | [256, 512] |

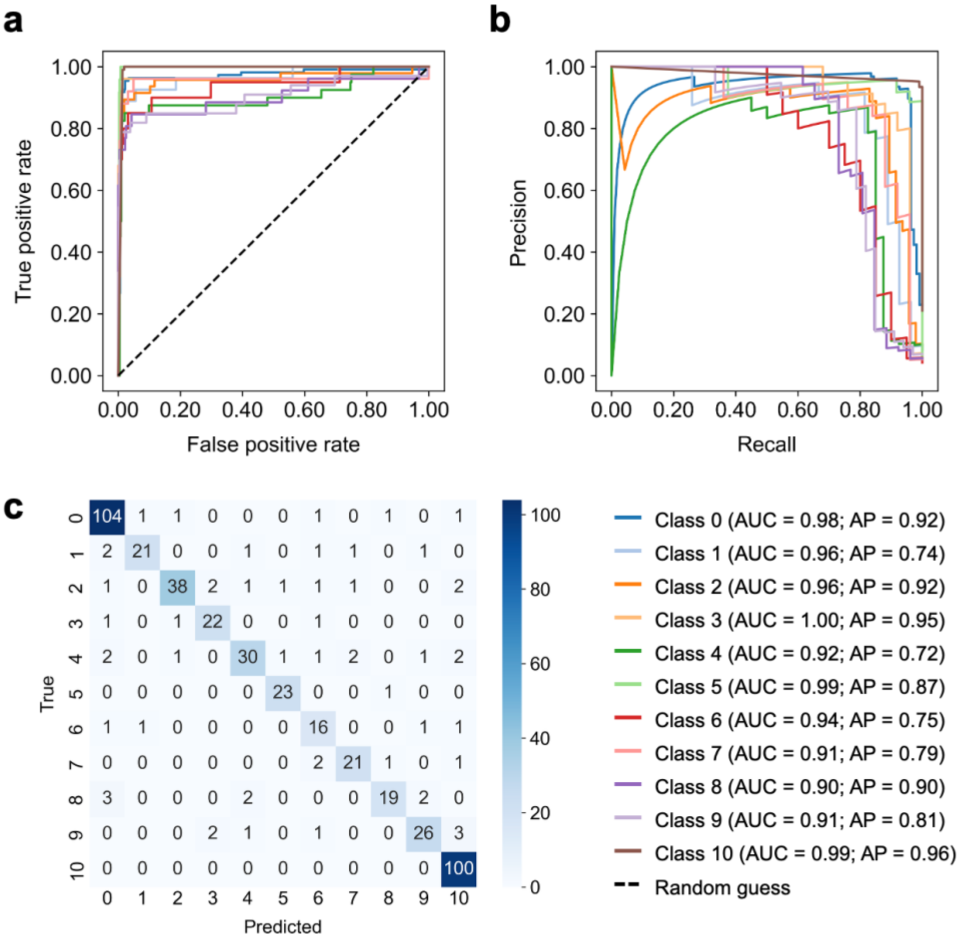


**Figure S4. Performance evaluation of the multilayer perceptron models.** (a) One-vs-rest receiver operating characteristic (ROC) curves for the multilayer perceptron model. (b) Precision-recall curves for the multilayer perceptron model. (c) Confusion matrix of the multilayer perceptron classifier for category-level prediction. Food categories (Classes 0-10) correspond to: alcoholic beverages, fats and oils, coffee/cocoa/tea, fish and seafood, fruit products, grain products, legumes/nuts/oilseeds/spices, meat products, seasonings and condiments, vegetable products, and others.

**Table S5. Summary of AI-generated aroma formulations by food category (classifier probability > 0.95), and their mean maximum similarity to existing aroma formulations.**

| Categories | Counts | Mean maximum similarity |
|---|---|---|
| alcoholic beverages | 997 | 0.73 |
| fats and oils | 904 | 0.60 |
| coffee, cocoa, tea | 823 | 0.55 |
| fish products | 951 | 0.73 |
| fruit products | 803 | 0.60 |
| grain products | 827 | 0.61 |
| legumes, nuts, oilseeds and spices | 712 | 0.53 |
| meat products | 730 | 0.68 |
| seasoning and condiments | 842 | 0.72 |
| vegetable products | 825 | 0.52 |
| others | 970 | 0.11 |

**Table S6. AI-generated aroma formulations (samples 1-3) and the positive control group used for sensory evaluation.**

| Samples | PubChem IDs | Compound names | Proportions |
|---|---|---|---|
| Sample 1 | 179 | 3-hydroxy-2-butanone | 0.1436 |
| | 264 | butanoic acid | 0.0159 |
| | 454 | 1-octanal | 0.0039 |
| | 7144 | creosol | 0.0243 |
| | 7165 | ethyl benzoate | 0.1281 |
| | 7342 | ethyl isobutyrate | 0.0077 |
| | 7762 | ethyl butanoate | 0.0600 |
| | 7797 | ethyl heptanoate | 0.0269 |
| | 7799 | ethyl octanoate | 0.0804 |
| | 7824 | methyl hexanoate | 0.0170 |
| | 8048 | ethyl decanoate | 0.0780 |
| | 12398 | heptadecane | 0.0137 |
| | 25611 | 2-methyl-3-heptanone | 0.0058 |
| | 31249 | diethyl succinate | 0.0060 |
| | 31253 | beta-Myrcene | 0.0072 |
| | 31265 | ethyl hexanoate | 0.1674 |
| | 88290 | 4-methyl-4-mercaptopentan-2-one | 0.0042 |
| | 440917 | D-limonene | 0.2101 |
| Sample 2 | 798 | indole | 0.0081 |
| | 7144 | creosol | 0.0163 |
| | 7165 | ethyl benzoate | 0.1437 |
| | 7762 | ethyl butanoate | 0.0439 |
| | 7797 | ethyl heptanoate | 0.0316 |
| | 7799 | ethyl octanoate | 0.0701 |
| | 7824 | methyl hexanoate | 0.0075 |
| | 8048 | ethyl decanoate | 0.0606 |
| | 12398 | heptadecane | 0.0237 |
| | 26808 | 2,3,5-trimethylpyrazine | 0.0038 |
| | 31249 | diethyl succinate | 0.0031 |
| | 31253 | beta-Myrcene | 0.0122 |
| | 31265 | ethyl hexanoate | 0.1233 |
| | 88290 | 4-methyl-4-mercaptopentan-2-one | 0.0042 |
| | 440917 | D-limonene | 0.4444 |
| | 853433 | isoeugenol | 0.0035 |

| | | | |
|---|---|---|---|
| Sample 3 | 179 | 3-hydroxy-2-butanone | 0.0324 |
| | 264 | butanoic acid | 0.0326 |
| | 454 | 1-octanal | 0.0552 |
| | 460 | guaiacol | 0.0211 |
| | 998 | phenylacetaldehyde | 0.1266 |
| | 3314 | eugenol | 0.0255 |
| | 6184 | 1-hexanal | 0.0919 |
| | 6549 | linalool | 0.0247 |
| | 8063 | pentanal | 0.0509 |
| | 8103 | 1-hexanol | 0.0438 |
| | 8130 | 1-heptanal | 0.0457 |
| | 11552 | 3-methyl butanal | 0.0935 |
| | 18827 | 1-octen-3-ol | 0.0836 |
| | 19309 | furaneol | 0.0205 |
| | 19310 | dimethyl trisulfide | 0.0216 |
| | 31265 | ethyl hexanoate | 0.0322 |
| | 31289 | nonanal | 0.1335 |
| | 637563 | anethole | 0.0216 |
| | 5283345 | (E)-2-decenal | 0.0428 |
| Control | 179 | 3-hydroxy-2-butanone | 0.0749 |
| | 323 | coumarin | 0.0119 |
| | 454 | 1-octanal | 0.0032 |
| | 1136 | sulfurol | 0.2165 |
| | 2758 | 1,8-cineole | 0.0844 |
| | 2879 | p-cresol | 0.0003 |
| | 3314 | eugenol | 0.0868 |
| | 6184 | 1-hexanal | 0.0045 |
| | 6549 | linalool | 0.0181 |
| | 6654 | alpha-pinene | 0.0053 |
| | 6920 | 2-acetylthiophene | 0.0045 |
| | 8063 | pentanal | 0.0011 |
| | 8103 | 1-hexanol | 0.0614 |
| | 8129 | 1-heptanol | 0.0027 |
| | 8130 | 1-heptanal | 0.0014 |
| | 8815 | estragole | 0.0023 |
| | 18635 | methional | 0.0013 |
| | 19309 | furaneol | 0.0020 |
| | 19310 | dimethyl trisulfide | 0.1732 |

| | | |
|---|---|---|
| 21059 | ethyl maltol | 0.0084 |
| 31244 | 4-methoxybenzaldehyde | 0.0110 |
| 31289 | nonanal | 0.0470 |
| 34286 | 2-methyl-3-furanthiol | 0.0024 |
| 61346 | 1-octen-3-one | 0.0023 |
| 83101 | 2,6-diethylpyrazine | 0.0020 |
| 520108 | 2-acetylthiazole | 0.0025 |
| 637563 | anethole | 0.1420 |
| 4564493 | 5-methyl-4-hydroxy-3(2H)-furanone | 0.0035 |
| 5282110 | cinnamyl acetate | 0.0088 |
| 5283345 | (E)-2-decenal | 0.0060 |
| 5283349 | 2,4-decadienal | 0.0025 |
| 5283356 | (E)-2-undecenal | 0.0058 |

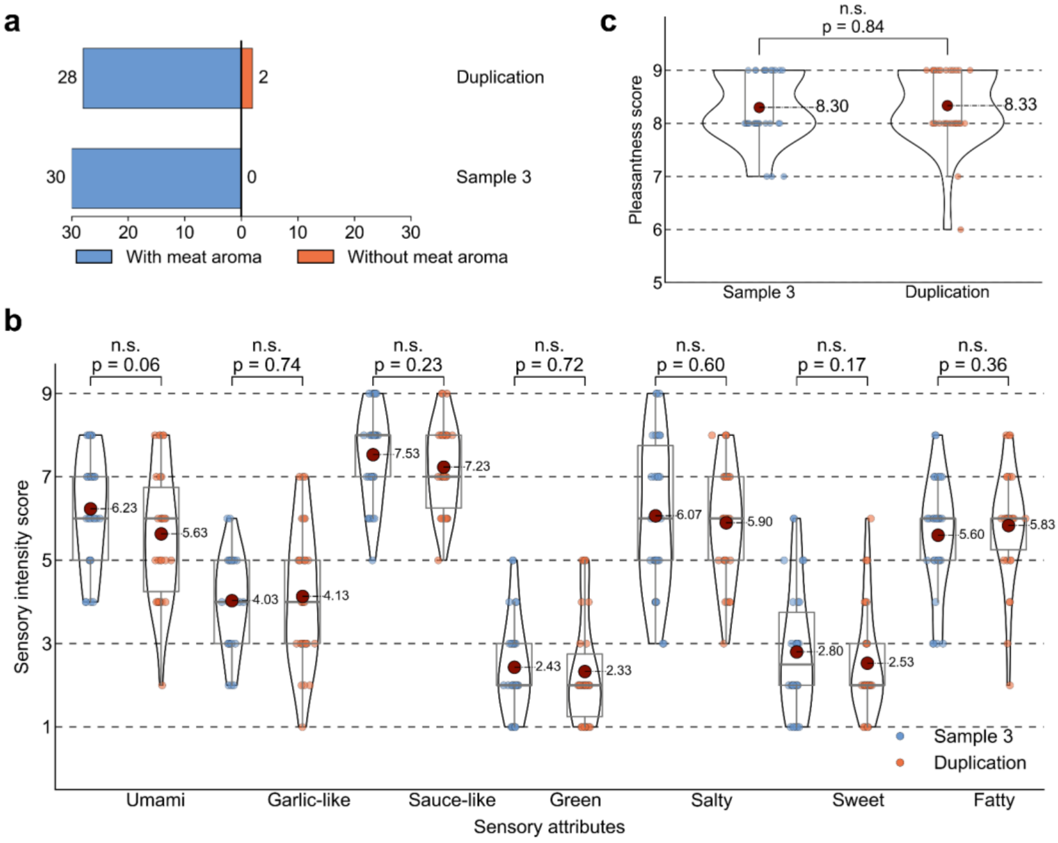


**Figure S5. Repeatability assessment of the sensory evaluation.** (a) Sensory panel judgments of the meat aroma of sample 3 and its duplicate. (b) Pleasantness scores for sample 3 and its duplicate, showing no significant differences. (c) Sensory intensity scores for sample 3 and its duplicate also showed no significant differences.

Note: In panels (b) and (c), the blue and orange scatter points denote individual ratings assigned by participants to each sample across different sensory dimensions. Grey box plots summarize the distributions for each group, displaying the first and third quartiles, the median, and the range of non-outlying values. Dark red points indicate the group means, and black curves represent the corresponding probability density distributions.

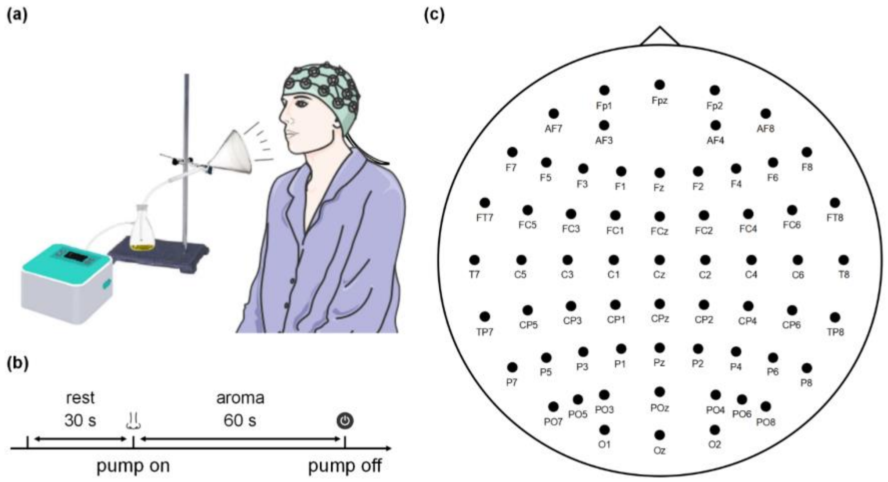


**Figure S6. Experimental setup for electroencephalography (EEG).** (a) Schematic of the aroma delivery system used during plant-based sausage tasting and EEG recordings. (b) Workflow of the EEG experiment. (c) Illustration of the 10-20 EEG electrode distribution.

**Table S7. Comparison of machine learning algorithms to predict aroma categories based on the proportion of KFOs as input features.**

| Machine learning algorithms | F1 scores ± standard deviation |
|---|---|
| Decision trees | 0.78 ± 0.02 |
| Linear discriminant analysis | 0.77 ± 0.01 |
| Logistic regression | 0.86 ± 0.01 |
| Multilayer perceptron | 0.87 ± 0.02 |
| Support vector machines | 0.86 ± 0.02 |
